# Comparative genomics reveals the origin of fungal hyphae and multicellularity

**DOI:** 10.1101/546531

**Authors:** Enikő Kiss, Botond Hegedüs, Torda Varga, Zsolt Merényi, Tamás Kószó, Balázs Bálint, Arun N. Prasanna, Krisztina Krizsán, Meritxell Riquelme, Norio Takeshita, László G. Nagy

## Abstract

Hyphae represent a hallmark structure of multicellular fungi with immense importance in their life cycle, including foraging for nutrients, reproduction, or virulence. Hypha morphogenesis has been the subject to intense interest, yet, the origins and genetic underpinning of the evolution of hyphae are hardly known. Using comparative genomics, we here show that the emergence of hyphae correlates with multiple types of genetic changes, including alterations of gene structure, gene family diversification as well as co-option and exaptation of ancient eukaryotic genes (e.g. phagocytosis-related genes). Half of the gene families involved in hypha morphogenesis have homologs in unicellular fungi and non-fungal eukaryotes and show little or no duplications coincident with the origin of multicellular hyphae. Considerable gene family diversification was observed only in transcriptional regulators and genes related to cell wall synthesis and modification. Despite losing 35-46% of their genes, yeasts retained significantly more multicellularity-related genes than expected by chance. We identified 414 gene families that evolved in a correlated fashion with hyphal multicellularity and may have contributed to its evolution. Contrary to most multicellular lineages, the origin of hyphae did not correlate with the expansion of gene families encoding kinases, receptors or adhesive proteins. Our analyses suggest that fungi took a unique route to multicellularity that involved limited gene family diversification and extensive co-option of ancient eukaryotic genes.

## Introduction

The evolution of multicellularity (MC) is considered one of the major transitions in the history of life^1^. It evolved in several pro-and eukaryote lineages^2–7^, each representing a unique solution to the challenges of multicellular organization^6^. Among the eukaryotes, two major modes for acquiring multicellularity are the clonal and aggregative routes^5,6,8–10^, which differ in how multi-celled precursors emerged by adhesion, cooperation, communication and functional diversification of cells^3,11,12^.

Fungi represent one of the three kingdoms where multicellular forms dominate among extant species^13^, yet, our knowledge on the evolutionary origin of multicellularity in this group is very incomplete. While most multicellular lineages can be recognized as either clonal or aggregative by comparisons to their unicellular relatives, fungal multicellularity has been recalcitrant to such categorization^6,14^. The thalli of fungi are made up of hyphae, thin, tubular structures that grow by apical extension to form a mycelium that explores and invades the substrate. Hyphal multicellularity has a number of unique properties compared to clonal and aggregative multicellularity, raising the possibility that its evolution follows markedly different principles^7^. First, hyphae might have evolved by the gradual elongation of substrate-anchoring rhizoids of early fungi^15–18^, through multinucleate intermediates, in contrast to clonal and aggregative lineages, where the first multi-celled clusters probably emerged via related cells sticking together (e.g. choanoflagellates^19^), or gathering to form a syncytial body (e.g. *Capsaspora*)^9^. Because early hyphae were uncompartmentarized, their evolution could have bypassed the need to resolve group conflicts and align the fitness of individual cells^7^. Alternatively, it is possible that conflicts are resolved at the level of individual nuclei^20^. Second, hyphae maximize foraging and nutrient assimilation efficiency and minimize competition for nutrients by a fractal-like growth mode^16,21,22^. This mode of origin differs from that of other multicellular lineages where selection for increased size possibly helped avoiding predation^2^. Hyphae might have also facilitated the transition of fungi to terrestrial life^23^ and confer immense medical relevance to pathogenic fungi^24^. Hyphae of extant fungi rarely stick to each other in vegetative mycelia and adhesion becomes key only in fruiting bodies^25–27^ - which, in terms of complexity level, resemble complex multicellular metazoans and plants^7,28^ - or in the attachment to host surfaces^29^. Thus, whereas in most multicellular lineages adhesion, cell-cell cooperation, communication and differentiation represent the main hurdles to the emergence of multicellular precursors^3,6,30–32^, fungi might have had different obstacles to overcome.

While the origins of hyphae are poorly known, information on the molecular and cellular basis of hyphal morphogenesis is abundant (for recent reviews see refs^33–38^), permitting evolutionary genomic analyses of the origins of hyphae. Hyphal morphogenesis builds on cell polarization networks^39^, the exo-and endocytotic machinery^40^, long range vesicle transport as well as fungal-specific traits such as cell wall synthesis and assembly^41^, and the selection of branching points and sites of septation^42^, among others. A key structure of hyphal growth is the Spitzenkörper^43^, which acts as a distribution center for vesicles transporting cell wall materials and various factors to the hyphal tip. The cytoplasmic microtubule network provides the connection between vesicle cargo through the ER and Golgi and the Spitzenkörper, from where vesicles move along actin microfilaments to the hyphal tip and secrete their content to deposit new cell wall components and provide surface expansion. Further key processes include the recycling of excess membrane in the subapical zone, the activation of cAMP pathways and mitogen activated protein kinase (MAPK) cascades and finally the transcriptional control of morphogenesis (for detailed reviews see refs^44–49^).

A complex hyphal thallus has been reported from a 407 million year old fossil Blastocladiomycota^51^, whereas Glomeromycotina-like hyphae and spores were preserved 460 million years ago^50,52^ indicating that hyphal growth dates back to at least the Ordovician. Most Dikarya and Mucoromycota grow true hyphae, whereas a significant diversity of forms is found in the early diverging Blastocladiomycota, Chytridiomycota and to a smaller extent the Zoopagomycota. The Chytridiomycota is dominated by unicellular forms that anchor themselves to the substrate by branched, root-like rhizoids^16,50^ which have been hypothesized as the precursors to hyphae^15,53^. An alternative hypothesis designates hypha-like connections in the thalli of polycentric chytrid fungi (e.g. *Physocladia*) as intermediates to true hyphae^17^. Like chytrids, most Blastocladiomycota form mono-or polycentric, unicellular thalli, although some species form wide, apically growing structures resembling true hyphae (e.g. *Allomyces*) or narrow exit tubes on zoosporangia (e.g. *Catenaria* spp.)^16,50,54^. In spite of these intermediate forms, the unicellular dominance in these phyla aligns well with a unicellular ancestry and potential convergent origins of hypha-like structures^17^.

Here we examine the evolution of hyphal multicellularity in fungi by reconstructing historical patterns of known hyphal morphogenesis genes as well as by systematic searches of fungal genomes for gene families whose evolution correlates with that of hyphae. We analyze the genomes of 4 plesiomorphically unicellular, 40 hyphal (one of which is ambiguous) and 14 secondarily simplified (yeast-like) fungi as well as 13 non-fungal relatives. Given the likely convergent origins of hyphae, we focus our analyses on multiple nodes of the fungal tree to where origin(s) of hyphal growth can be localized with confidence. Our analyses reveal a deep eukaryotic origin of most morphogenesis-related families, limited gene family diversification in correlation with the emergence of hyphal MC and that secondarily simplified yeast-like fungi retained most of the genes for multicellular growth.

## Results and discussion

### Hyphae evolved in early fungal ancestors

To understand the origin of hyphae, we constructed a species phylogeny representing 71 species (Supplementary Table 1) by maximum likelihood analysis of a supermatrix of 595 single-copy orthologs (175,535 characters Fig. 1a).

**Fig. 1.**
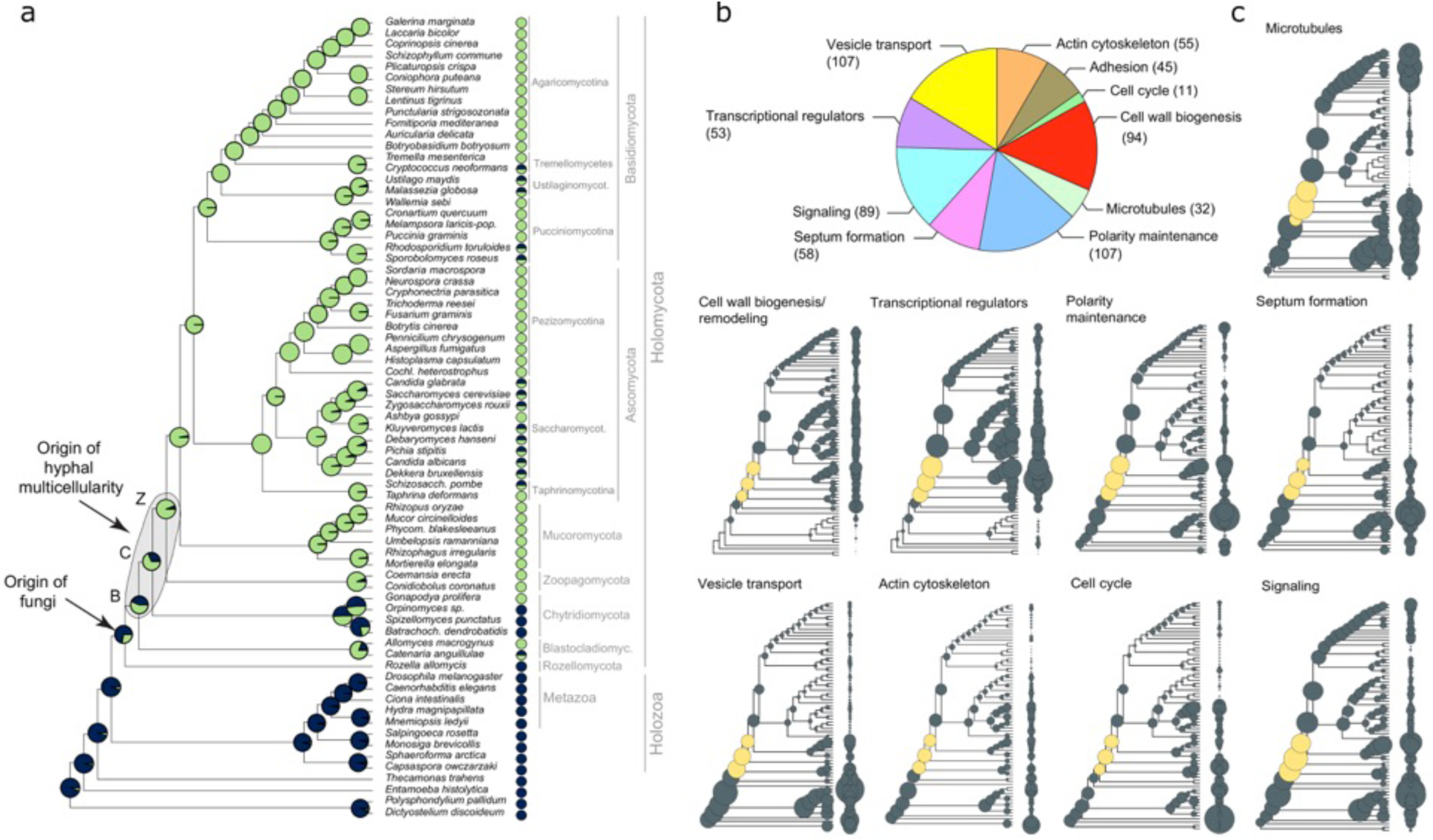
The evolution of hyphal multicellularity and underlying genes in fungi. (a) Phylogenetic relationships among 71 species analyzed in this study. Pie charts at nodes show the proportional likelihoods of hyphal (green) and non-hyphal (dark blue) ancestral states reconstructed using Bayesian MCMC. Character state coding of extant species used in ancestral state reconstructions is shown next to species names. BCZ nodes: origin of hyphal growth could be assigned with confidence are highlighted (note the uncertainty imposed by filamentous Blastocladiomycota). (b) the distribution of literature-collected hypha morphogenesis genes among 10 main functional categories. (c) Ancestral reconstructions of gene copy number in 9 main hypha morphogenesis-related categories of genes (see Fig. 5c for adhesion). Bubble size is proportional to reconstructed ancestral gene copy number. BCZ nodes are shown in yellow.

Our species phylogeny recapitulates recent genome-based phylogenies of fungi^55–58^, with the Rozellomycota, Blastocladiomycota and the Chytridiomycota splitting first, second and third off of the beckbone, respectively (ML bootstrap: 100%). We performed ancestral character state reconstruction using Bayesian MCMC to identify the putative origin of hyphal growth. This supported an emergence of hyphae from unicellular ancestors in basal fungi. The distribution of posterior probability values indicated three nodes as the most likely origins of hyphal multicellularity, which represent the split of Blastocladiomycota, Chytridiomycota and Zoopagomycota lineages, referred hereafter to as BCZ nodes. The posterior probability for the hyphal state started to rise in the most recent common ancestor (MRCA) of the Blastocladiomycota and higher fungi (PP: 0.53, Fig. 1a) and increased to 0.68 and 0.92 in the next two nodes along the backbone of the tree. This suggests that hyphae evolved either in one of the BCZ nodes or it may have been a gradual process unfolding in these three nodes. This uncertainty likely reflects diverse hypha-like morphologies in the Blastocladio-and Chytridiomycota and is consistent with the convergent origins of hypha-like morphologies^7,17,18^. To account for this uncertainty, we focus on BCZ nodes in subsequent analyses of hypha morphogenesis genes.

### The evolution of hypha morphogenesis genes

Our survey of the literature for hyphal multicellularity-related genes yielded 651 genes (from 519 publications), mostly from well-studied model systems such as *A. fumigatus, A. nidulans, N. crassa, S. cerevisiae* and *C. albicans* (Supplementary Table 2). We categorized genes according to the broader function they fulfill in hyphal growth into nine functional groups: actin cytoskeleton regulation, polarity maintenance, cell wall biogenesis/remodelling, septation (including septal plugging), signaling, transcriptional regulation, vesicle transport, microtubule-based transport and cell cycle regulation. The categories “polarity maintenance” and “vesicle transport” contained the largest number of genes (107 in each), whereas “cell cycle regulation” contained the fewest (11) (Fig.1b). The collected genes grouped into 362 families by Markov clustering of the a 71-genome dataset.

Reconstructions of gene duplication/loss histories for nine functional categories of hypha morphogenesis gene families are shown on Fig. 1c. A general pattern that emerges from these is that the origin of many gene families (181 families, 50%) predate that of hyphal MC (Fig. 2, Supplementary Fig. 1), indicating that fungi have co-opted several conserved eukaryotic functionalities for hyphal growth. A significant proportion of multicellularity-related gene families (164 families, 45.3%) emerged after the origin of hyphal MC, indicating lineage-and species-specific genetic innovations. Only 17 families (4.7%) originated in BCZ nodes and were conserved thereafter (Table 1), providing potential candidates that shaped the evolution of hyphal MC. These include two families of transcriptional regulators (encoding StuA and MedA proteins in *A. fumigatus*)^59,60^, six related to cell wall biogenesis, three to actin cytoskeleton regulation, three to polarity maintenance, two families involved in signaling and one involved in cell cycle regulation. One of the families contains the *S. cerevisiae* Pan1, an endocytic adaptor protein at the plasma membrane. Pan1 triggers the recruitment of the Arp2/3 complex to the site of endocytosis, which is necessary for the recycling of excess membrane in the subapical region during hyphal growth^40^. Another example is the polarisome component BNI-1 from *N. crassa*. Knockout studies showed that it mediates actin cable assembly in filamentous fungi and has a role in diverse morphogenesis-related processes^61^. Other proteins involved in establishing cell polarity are the Bem1 actin cytoskeleton reorganizing factor^62,63^ and the Rax1, associated with bipolar budding in *S. cerevisiae*^64^.

**Fig. 2.**
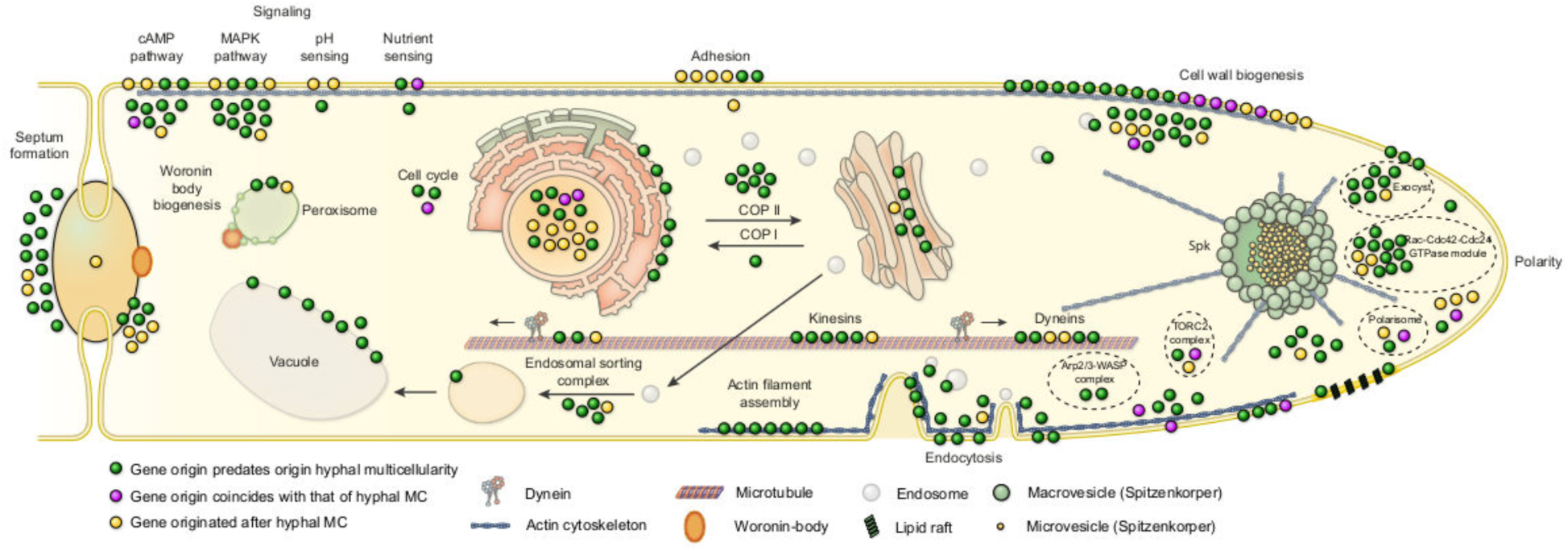
Phylogenetic age distribution of hypha morphogenesis genes. Schematic outline of terminal hyphal cell is shown with genes marked by dots and colored by phylogenetic age. Genes whose origin predates that of hyphal multicellularity (green, 72,2%) dominate the hyphal morphogenetic machinery, followed by genes that originated after hyphal MC (yellow, 21,2%) and genes whose origin approximately coincides with that of hyphae (purple, 6,6%). Data based on only *A.fumigatus* orthologs. See also Supplementary Figure 1 for gene names.

**Table 1:** List of the 17 gene families whose emergence shows correlation with the evolution of hyphal MC based on COMPARE analysis.

| Emergence of gene family | *A. fumigatus* ortholog | *S. cerevisiae* ortholog | functional category | cluster ID |
| --- | --- | --- | --- | --- |
| Ica of Dikarya, Mucoromycota, Zoopagomycota, Chytridiomycota, Blastocladiomycota | Afu7g03870 | PAN1/YIR006C | actin cytoskeleton | 4903 |
|  | crh3/Afu3g09250 | UTR2/YEL040W | cell wall biogenesis | 374 |
|  | gel7/Afu6g12410 | GAS1/YMR307W | cell wall biogenesis | 435 |
|  | Afu6g04940 | BNR1/YIL159W | polarity establishment | 3689 |
|  | Afu4g04120 | BEM1/YBR200W | polarity establishment | 2482 |
|  | stuA/Afu2g07900 | PHD1/YKL043W | transcriptional regulation | 2479 |
|  | medA/Afu2g13260 | NA | transcriptional regulation | 8521 |
| Ica of Dikarya, Mucoromycota, Zoopagomycota, Chytridiomycota | Afu6g07910 | SLM1/YIL105C | actin cytoskeleton | 1561 |
|  | Afu8g04520 | SLA1/YBL007C | actin cytoskeleton | 5953 |
|  | Afu4g06130 | WHI2/YOR043W | cell cycle regulation | 2915 |
|  | Afu4g00620 | DFG5/YMR238W | cell wall biogenesis | 843 |
|  | Afu8g02320 | NA | cell wall biogenesis | 3398 |
|  | chsD/Afu1g12600 | NA | cell wall biogenesis | 9065 |
|  | rgsB/Afu4g12640 | RAX1/YOR301W | polarity establishment | 2900 |
|  | Afu2g08800 | SSY1/YDR160W | signaling | 49 |
|  | ricA/Afu4g08820 | NA | signaling | 5950 |
| Ica of Dikarya, Mucoromycota, Zoopagomycota | kre6/Afu2g11870 | KRE6/YPR159W | cell wall biogenesis | 293 |

Gene families related to septation, polarity maintenance, cell cycle control, vesicle transport and microtubule-based transport are generally more diverse in animals, non-fungal eukaryotes and their ancestors than in fungi, suggesting that despite the key role of these families in hyphal MC, they evolved primarily by gene loss in fungi (Fig 1c). Cell wall synthesis and remodeling as well as transcription regulation related families, on the other hand, show expansions in MC fungi, suggesting that the diversification of these gene families could have played roles in the evolution of hyphal MC (Fig 1c).

Ninety-three (25.7%) of the 362 hyphal morphogenesis-related gene families showed duplications in BCZ nodes (Supplementary Table 3). Enrichment analysis of gene duplications revealed no individual gene families with significantly increased number of duplications (Benjamini-Hochberg corrected P<0.05, Fisher’s exact test) in BCZ nodes, relative to the rest of the tree (Supplementary Table 4). The same analysis on the 9 functional groups showed significantly increased numbers of duplications in the cell wall biogenesis and transcriptional regulation categories. These analyses suggest that the evolution of hyphal growth in BCZ nodes did not generally coincide with a period of extensive gene duplication except in cell wall biogenesis and transcriptional regulation-related gene families. Collectively, reconstructions of gene family evolution revealed the lack of a major burst of gene family origin or that of duplications coincident with the evolution of hyphal MC.

We analyzed whether changes in basic structural properties of genes show a correlation with the evolution of hyphal MC. Significant differences (P < 0.05) were observed in gene, CDS and intron lengths between unicellular and multicellular fungi (Supplementary Fig. 2, Supplementary Table 5). CDS lengths of septation and polarity maintenance genes were significantly higher in multicellular than in unicellular fungi (P=0.0012-0.00017, Supplementary Fig. 2). An opposite pattern was observed in intron lengths, which were on average longer in unicellular fungi in actin cytoskeleton, cell wall biogenesis, polarity maintenance, septation and vesicle transport related genes. In actin cytoskeleton-related genes the CDS length and intron length showed the same pattern: both of them were longer in multicellular species. Gene and CDS lengths were significantly longer in unicellular than in multicellular fungi in genes encoding adhesion and microtubule-based transport proteins. On the other hand, interestingly, no significant changes in gene structure were detected in cell wall biogenesis and transcriptional regulation-related genes, the two categories that displayed significant gene family diversification in early filamentous fungi.

### Phagocytosis is lost, but phagocytotic genes are retained by fungi

Our sets of hypha morphogenesis genes included several entries associated with phagocytosis in non-fungal eukaryotes. This is surprising given that phagocytosis is not known in fungi and their rigid cell wall forms a physical barrier to it. We therefore examined the fate of phagocytosis genes in filamentous fungi based on the phagocytotic machinery of *D. discoideum*^65,66^ and other eukaryotes^67^. Filamentous fungi have retained several phagocytotic gene families but lost others (Fig. 3). For example, members of the Arp2/3 complex, which nucleates actin filaments and triggers actin cytoskeleton rearrangements^68^ is conserved in filamentous fungi and is involved in hyphal growth^69^. Engulfment and cell motility genes (ELMO1/2) are found in all filamentous fungi, but are convergently lost in budding and fission yeasts as well as in *C. neoformans, M. globosa* and *W. sebi*, all of which have reduced capacities for hyphal growth. The DOCK (dedicator of cytokinesis) protein family, which interacts with ELMO proteins, is represented as a single gene copy in filamentous fungi and yeasts. This family contains orthologs of *S. cerevisiae* DCK1 (a homolog of human DOCK1), which has been shown to influence hypha morphogenesis^70^. Of the broader Wiskott-Aldrich syndrome family of proteins, which reorganize the actin cytoskeleton during phagocytosis, the WASP family is conserved across fungi, the WAVE family is only represented in early diverging fungi and non-fungal eukaryotes, whereas the WASH family has been lost in fungi, with homologs detected only in non-fungal eukaryotes, consistent with recent reports^71,72^. These patterns reveal the conservation of several phagocytotic genes in fungi, despite the loss of phagocytosis and the evolution of rigid cell walls and osmotrophy. Previous functional analyses have shown that several of the retained genes are involved in hypha morphogenesis in fungi, indicating that members of the phagocytic machinery were probably exapted for hyphal multicellularity in ancient fungi.

**Fig. 3.**
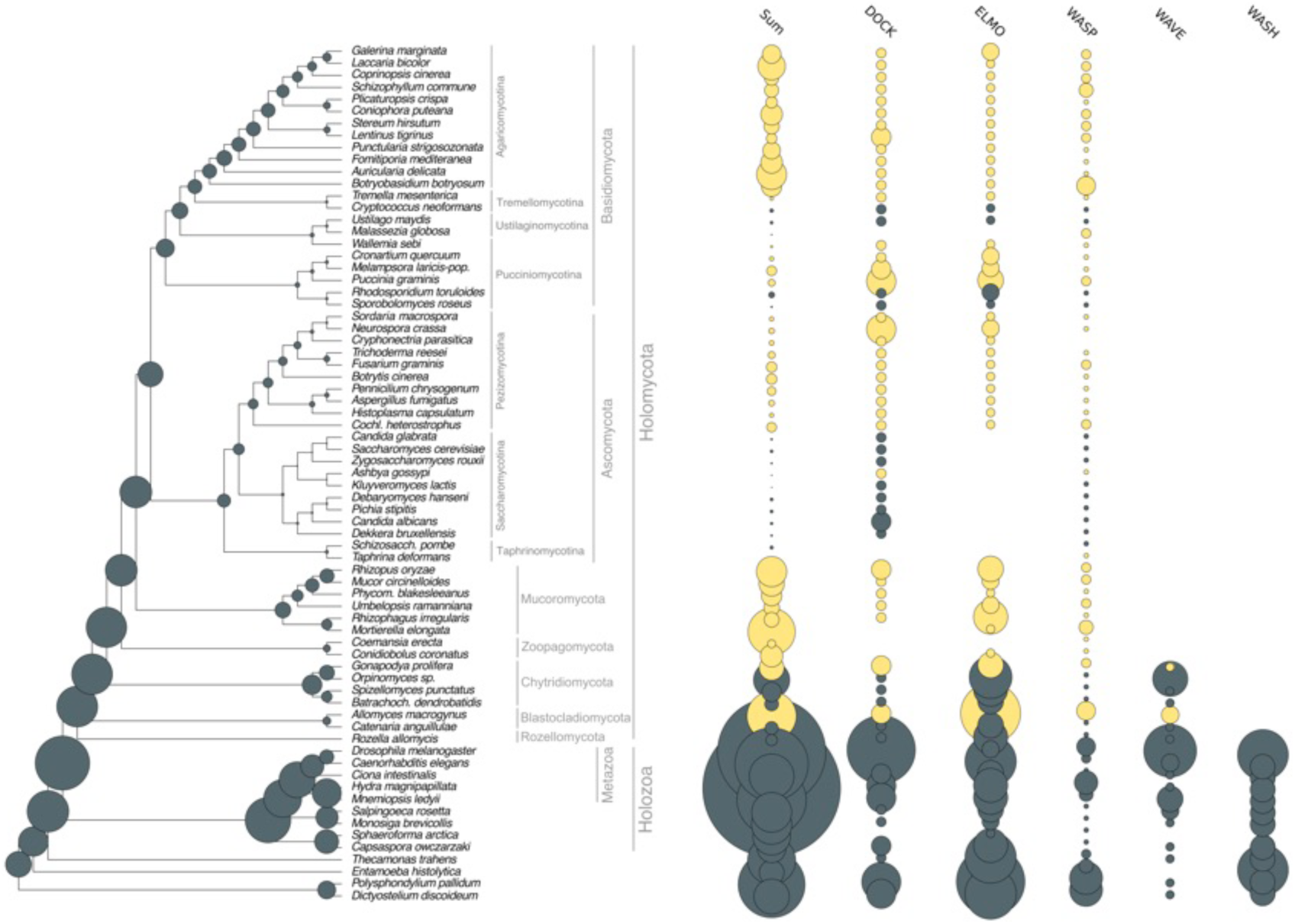
Evolutionary dynamics of phagocytosis-related gene families. Several phagocytic gene families retained in filamentous fungi (DOCK, ELMO, WASP). WAVE family retained only in early fungi (Blastocladiomycota and Chytridiomycota), WASH family is represented only in non-fungal eukaryotes. Bubble size is proportional to ancestral and extant gene copy number. Copy numbers of filamentous fungi are labelled with yellow.

### Genome-wide screen for correlated evolution between multicellularity and gene family expansion

To identify further gene families with a potential connection to hyphal MC, we systematically searched for families that show substantial dynamics in BCZ nodes. We reasoned that gene families underlying hyphal MC should originate or diversify in BCZ nodes and be conserved in descendent filamentous fungi. Searching for gene families fitting these criteria yielded 414 families (ANOVA, p<0.05, Supplementary Table 6), 114 of which originated in BCZ nodes, while the others showed duplication rates that exceeded the expectation derived from genome-wide collection of gene families (Fig. 4). These included several known morphogenetic families (e.g. Bgt3, RgsB and Gel2 of *A. fumigatus*, Bem1 and Rax1 of *S. cerevisiae*), genes involved in actin cytoskeleton and cell wall assembly, mating, pheromone response (GpaA of *A. fumigatus*), sporulation and transporters, among others (Supplementary Table 6). Several of the identified families contain genes with reported growth defects in *A. fumigatus* or *S. cerevisiae*, indicating that our searches recovered genes relevant for hyphal MC. For example, Rax proteins are major regulators of cellular morphogenesis and are involved in bud site selection in budding yeasts^73,74^, polarized growth in *S. pombe*^75^ and polarity maintenance in filamentous fungi^76^. The finding that these families originated in BCZ nodes makes them candidates for being key contributors to the evolution of hyphal MC. We further detected a fungal-specific cluster of tropomyosins (TPM1 in *S. cerevisiae*), which maps to the MRCA of Blastocladiomycota and other fungi and comprises genes involved in polarized growth and the stabilization of actin microfilaments. The family containing *S. pombe* Dip1 homologs (Afu6g12370 in *A. fumigatus*) emerged in the node uniting Chytridiomycota with higher fungi and contains a single gene per species afterwards, except an expansion in WGD Mucoromycota^77^ and losses in the Saccharomycotina. In *S. pombe*, Dip1 activates the Arp2/3 complex without preexisting actin cables^78,79^ and thus initiates cortical actin patch assembly and endocytosis. Because it does not require pre-existing actin cables, it mediates actin cytoskeleton regulation through a mechanism that seems to be specific to multicellular fungi. Finally, we detected the family containing *S. cerevisiae* Dpp1 homologs, which shows a significant expansion (4 duplications) in BCZ nodes. This family regulates morphogenetic transitions in dimorphic fungi through the synthesis of the fungal signal molecule farnesol^80^, which prompts us to speculate that it might have contributed to the elaboration of farnesol-based communication in fungi.

**Fig. 4.**
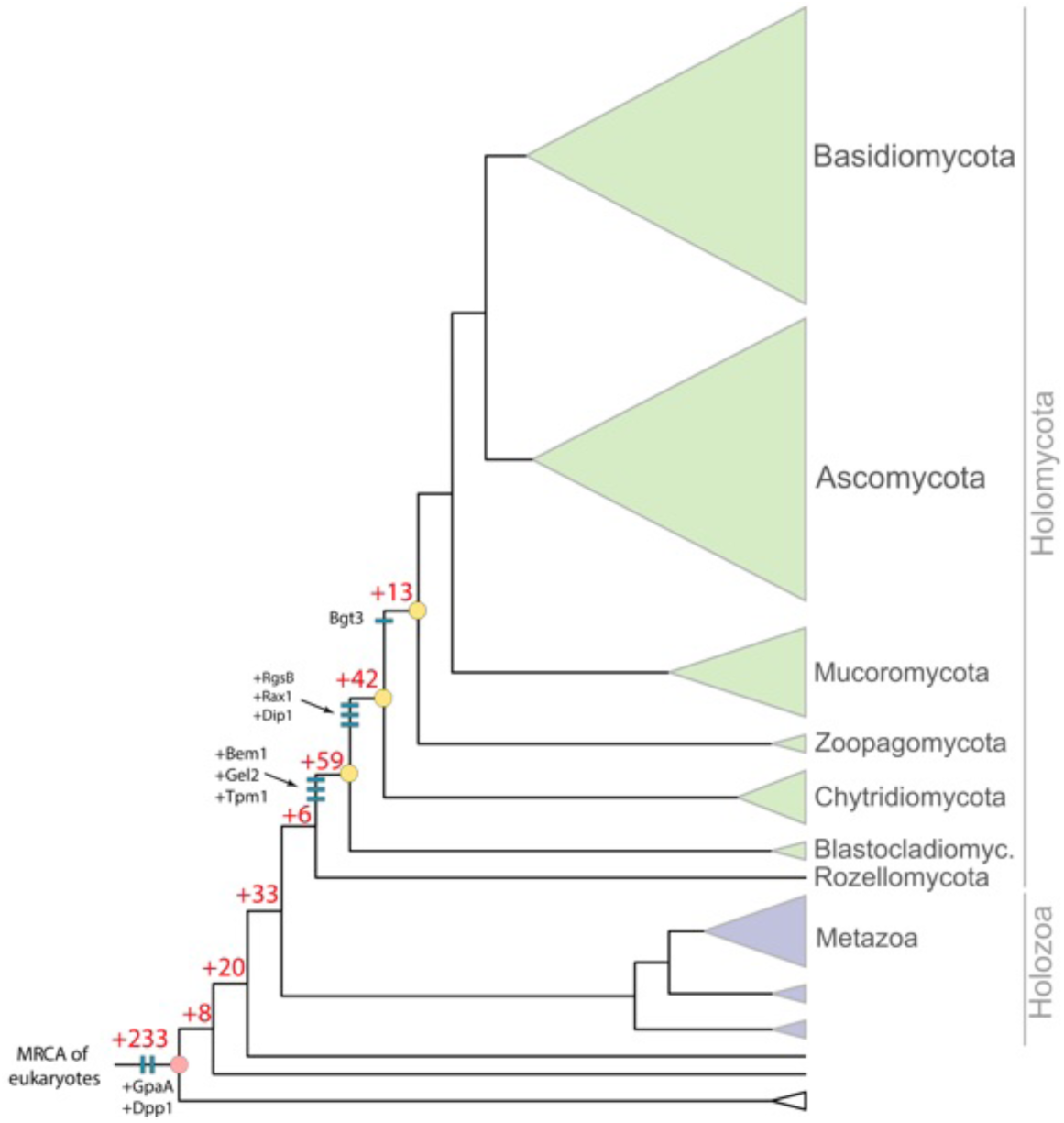
Origin of 414 gene families potentially related to the evolution of hyphal MC, identified by ANOVA (p<0.05). 114 families originated in BCZ nodes, including known morphogenesis related proteins (e.g. Bgt3, RgsB, Gel2 of *A. fumigatus*, Rax1, Bem1, Tpm1 and Dpp1 from *S. cerevisiae*, Dip1 from *S. pombe*) labelled as blue bars. Red numbers at branches represent the number of gene families originated at the backbone of the species tree.

Because the emergence of hyphal multicellularity overlaps significantly with that of other fungal traits, it is challenging to unequivocally separate signals conferred by these traits from those of hyphal MC. It is conceivable that a portion of the 414 gene families (Supplementary Table 6) were detected because of signals conferred by phylogenetically co-distributed traits, not necessarily multicellularity itself (see Beaulieu 2016^81^ for a conceptually analogous problem in analyzing taxonomic diversification). One such trait could be osmotrophy, the feeding mechanism of fungi to absorb soluble goods generated by extracellular enzyme complexes^82^. We detected 20 gene families that showed strong correlation with hyphal MC and were annotated as various transporters; such families could hypothetically be related to osmotrophy. Further, among the 414 detected families, there were 84 that are currently functionally uncharacterized and thus it is impossible to speculate about their role in hyphal MC. Collectively, these families indicate that there are plenty of fungal genes that evolved in concert with hyphal MC and that await functional characterization to establish links to hyphal MC or other fungal functions.

### The evolution of kinase, receptor and adhesive repertoires do not correlate with hyphal multicellularity

The increased sophistication of cell-cell communication and adhesion pathways often correlates with expanded repertoires of genes encoding kinases, receptors and adhesive proteins^83–85^. We therefore, examined Ser/Thr kinase (954 clusters), hybrid histidine kinase (96 clusters), receptor (183 clusters) and adhesion (23 clusters) genes, focusing on the comparison of unicellular and filamentous fungi. Copy numbers across the 954 identified Ser/Thr kinase clusters were similar in unicellular and simple multicellular fungi, with higher kinase diversity found in complex multicellular Basidiomycota (as reported by Krizsan 2018)^86^ and in *Rhizophagus irregularis* (Fig. 5a). We inferred net contractions in BCZ nodes, from 572 to 529 reconstructed ancestral kinases (81 duplications, 124 losses, Fig. 5a). Nevertheless, kinase families that duplicated here include all 3 MAPK pathways in fungi, the mating pheromone, cell wall integrity (*fus3, kss1, kdx1* from *S. cerevisiae, mpkB* from *A. fumigatus*) and osmoregulatory pathways (*hog1, ssk2, ssk22* from *S. cerevisiae, sakA, sskB* from *A. fumigatus*), all of which indirectly regulate hyphal growth^46–48^.

**Fig. 5.**
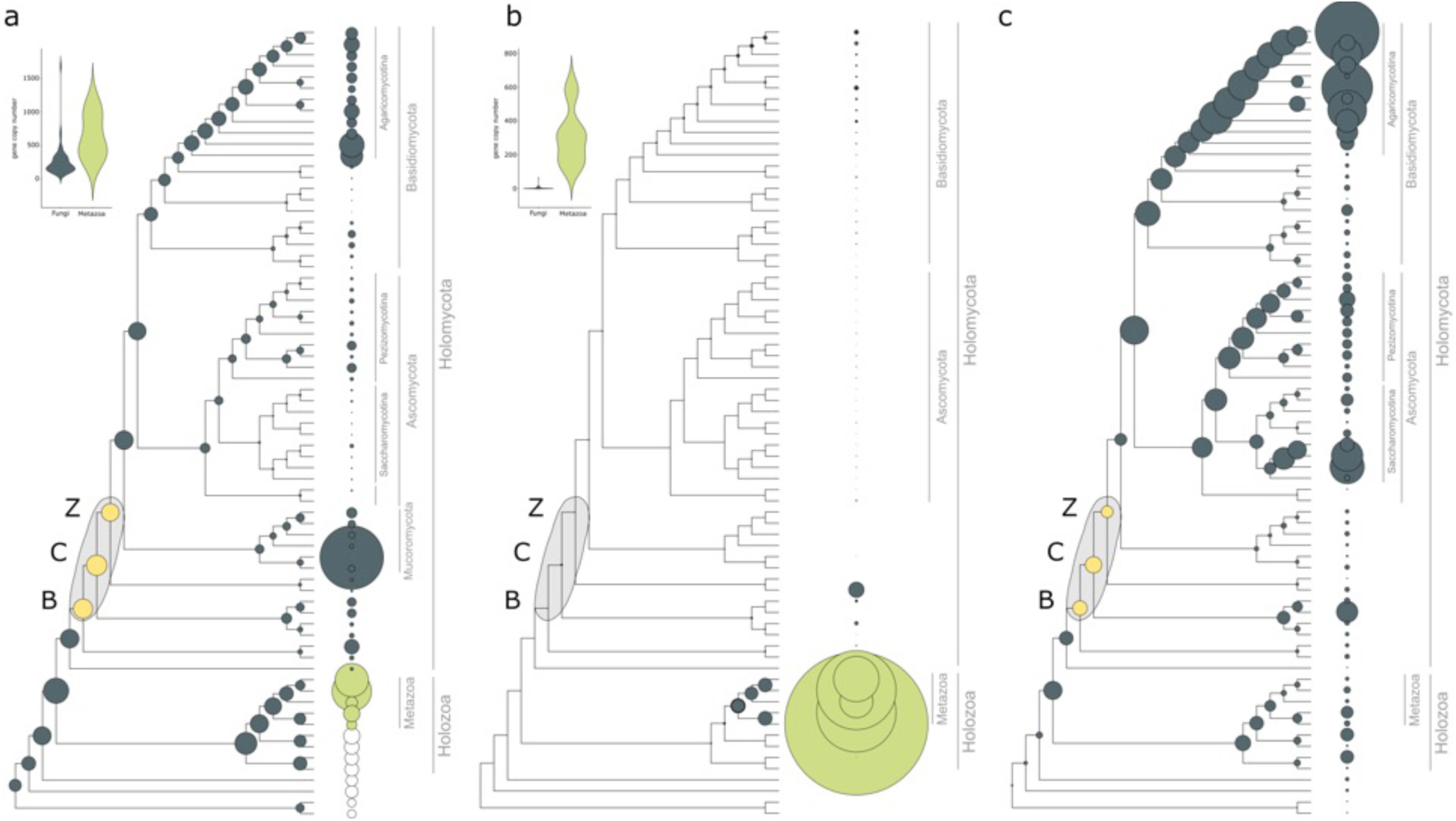
Gene family evolution of Serine-Threonine kinases (a), transmembrane receptors (b) and adhesion-related genes (c). BCZ nodes (yellow) represent the putative origin of hyphal MC. Bubble size across the tree is proportional to reconstructed ancestral gene copy number and copy number of extant species (shown right to the tree). Violin plots for kinases (a) and receptors (b) show copy number distribution of gene families in multicellular fungi (grey) and metazoans (green).

Overall, fungi had fewer Ser/Thr kinases (mean 257) than metazoans (mean 643). However, the higher kinase diversity of metazoans seems to be a result of an early expansion in the MRCA of Holomycota and Holozoa (Fig. 5a). While signal transduction requirements of metazoan MC have been mostly discussed in the context of receptor tyrosine kinases, we found no evidence for domain architectures typical of receptor tyrosine kinases in fungi. The only group resembling receptor kinases is hybrid histidine kinases (HK), that comprise proteins with a sensor domain, a histidine kinase domain and a C-terminal receiver domain that acts as a response regulator. We inferred an expansion (24 duplications, 10 losses) of HKs in the MRCA of the Chytridiomycota and other fungi, including class III and X HKs, which are linked to morphogenesis^87,88^. Another wave of HK expansion was inferred in the MRCA of Mucoromycota and Dikarya with 11 duplications and 4 losses (Supplementary Fig. 3).

In G-protein coupled receptors (GPCRs), an even more extreme difference was observed between fungi and metazoans (Fig. 5b). A large receptor expansion was observed in the latter, which resulted in 135-583 genes in extant animals. Out of the 183 analyzed GPCR families, only 19 were found in fungi, and only one of them was generally conserved across fungal species. This family contains STE3 and STE3-like a-factor mating pheromone receptors, involved in pheromone-dependent signal transduction, cellular conjugation and cell fusion.

Adhesive cell surface proteins are key mediators of the transition to MC in colonial and aggregative lineages^3,5,6^, which is reflected in their higher copy numbers in multicellular organisms^89^. We identified 45 families of putative adhesion-related proteins in fungi, including adhesins, flocculins, hydrophobins, various lectins and glycosylphosphatidylinositol (GPI)-anchored cell wall proteins. Our reconstructions of the evolution of these families (Fig. 5c) revealed no expansion but a small contraction (from 17 to 14 copies) in BCZ nodes. Expansions were inferred in the Agaricomycotina and in Saccharomycotina yeasts (*C. albicans, P. stipitis, hansenii*). The expansion in the Agaricomycotina was driven by class1 hydrophobins and homologs of the *C. neoformans* Cfl1 (an adhesive protein with roles in signaling and morphogenesis regulation)^90^. Diversification of these families correlates with the evolution of fruiting bodies and probably reflects the emergence of complex multicellularity^7^. The higher copy numbers in yeast species relate to several yeast-specific adhesin and lectin-like cell wall proteins that contain experimentally characterized adhesive proteins of human pathogenic fungi (e.g. *Candida* spp.)^91,92^.

Taken together, the evolution of kinase, receptor and adhesive protein repertoires highlights an important difference between fungi and other multicellular lineages. We observed no significant expansion of such families in filamentous fungi, whereas kinase and adhesion related genes expanded in complex multicellular Agaricomycotina. This might be explained by the two-step nature of the evolution of complex MC in fungi^7,93^ that proceeds through an intermediate complexity level, hyphal MC, as opposed to metazoans, where complex MC evolved in a more direct way^14^.

### Yeasts retain most genes required for hyphal morphogenesis

Yeasts are secondarily simplified organisms with reduced ability to form hyphae and that spend most of their life cycle as unicells^16,18,53,94^. Our ancestral character state reconstructions imply that yeasts derived from filamentous ancestors (Fig. 1a), and thus they represent a classic example of reduced complexity. They were hypothesized to have lost MC^95^, even though rudimentary forms of hyphal growth (termed pseudohyphae) exist in most species. We scrutinized the fate of MC-related genes in five predominantly yeast-like lineages^94^, the Saccharomycotina, Taphrinomycotina, Pucciniomycotina, Ustilaginomycotina and Tremellomycotina. Because yeast genomes have undergone extreme streamlining during evolution, we evaluated gene loss among MC-related genes in comparison to genome-wide figures of gene loss.

Yeast species generally have fewer MC-related genes and reconstructions indicate more losses than duplications along branches of yeast ancestors (Fig. 6). However, when we corrected for genome-wide reductions in gene number, we found that hyphal morphogenesis genes are underrepresented among lost genes compared to other functions (Fig. 6a, Supplementary Table 7). Most groups of hyphal morphogenesis genes are significantly depleted (P<0.05, Fisher’s exact test) among gene losses in yeast clades (except the Ustilaginomycotina). We recovered only 6 cases where losses of MC-related genes were significantly overrepresented (P<0.05, Fisher’s exact test, Supplementary Table 7), in proteins related to adhesion and microtubule-based transport. Higher than expected loss rates in such proteins were observed in the Taphrinomycotina, Pucciniomycotina, Tremellomycotina and the Saccharomycotina, suggesting that microtubule-based transport and adhesion-related functions become dispensable for these yeasts clades. Losses in ‘microtubule system’ are particularly interesting from the perspective of long-range transport of vesicles and nuclei along hyphae. Microtubule system-related genes with reduced repertoires in yeasts include gamma-tubulin complex proteins, kinesins, dynactin, dynein heavy chain (nudA), and the dynactin linking protein ro-2^96^. Of the five yeast-like clades, the budding and fission yeast lineages show the most gene losses, consistent with the strongest reduction of hyphal growth abilities in these clades. A similar pattern was found for NADPH oxidases, which is consistent with previous reports: concurrent losses in all yeast-like clades, with complete loss of the family in the Saccharomycotina, Ustilaginomycotina, in *S. pombe* and *C. neoformans* (Supplementary Fig. 4).

**Fig. 6.**
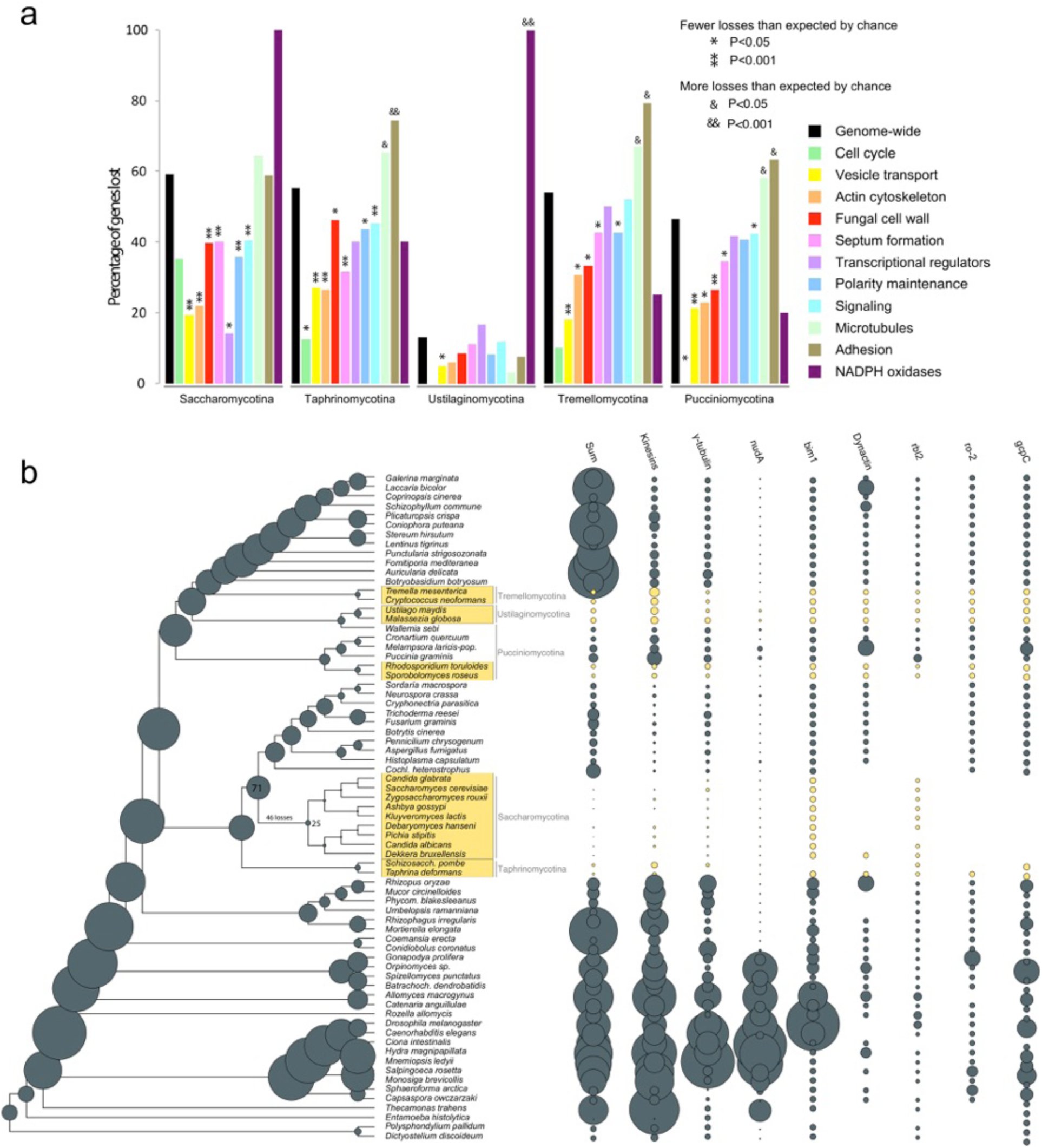
Secondarily simplified yeast-like fungi retain genes for hyphal MC. (a) the percentages of lost genes in main morphogenesis-related categories. Percentages were calculated relative to ancestral copy numbers inferred in the node preceding the origin of 5 yeast-like clades (Saccharomycotina, Taphrinomycotina, Pucciniomycotina, Ustilaginomycotina and Tremellomycotina). Significance of the enrichment of gene losses in each category relative to genome-wide figures of gene loss were determined by Fisher’s exact test and is shown above bars. (b) ancestral gene copy number reconstruction of microtubule-based transport genes along the fungal phylogeny. Secondarily simplified (yeast-like) clades are highlighted in yellow. Bubble size proportional to reconstructed ancestral and extant gene copy number across 19 gene families. Copy number distribution of each gene family is shown right to the tree.

Altogether 54-65% of MC-related genes were retained in yeast genomes (Supplementary Table 7). For example, hardly any reductions are observed in PI4,5P2 binding proteins (Slm1, Slm2 in *S. cerevisiae*^97^), which are involved in the regulation of actin cytoskeleton organization and endosomal transport, or in class I myosins (myoA in *A. nidulans*^98^, myo5 in *S. cerevisiae*^99^) which are localized to the site of polarized growth and involved in secretion and cell wall biogenesis.

Collectively, these data suggest that hyphal morphogenesis genes in general are dispensable for yeasts to a smaller extent than genes with other functions. This is consistent with most yeast-like fungi being able to switch to hyphal or pseudohyphal growth under certain conditions. The fact that hyphal morphogenesis genes are not statistically significantly enriched among losses compared to genome-wide expectation, however, is not in conflict with a reduction of multicellular abilities in yeast-like fungi. These gene losses do indicate reductions in hyphal growth, although this cutback is smaller compared to other functions in the genome. This in turn suggests, that the ability for multicellular growth is among the functions preferentially retained by yeast-like fungi.

## Conclusions

We analyzed the genetic underpinnings of the evolution of fungal hyphae. Hyphae are among the most enigmatic fungal structures with a unique multicellular organization, yet their evolutionary origins remained poorly explored. Our ancestral character state reconstructions localized the origin of hyphae to three nodes around the split of Blastocladio-, Chytridio-and Zoopagomycota (BCZ nodes), consistent with previous studies^17^ and potential convergence of hypha-like structures in these phyla.

To understand how the underlying genetics evolved, we identified 362 gene families with known relevance to hyphal morphogenesis (e.g. from knockout studies) and predicted a link to hyphal MC for another 414 families using comparative genomics of 71 species. The evolutionary dynamics of these families shows a mixed picture. A large proportion of families are conserved in all sampled eukaryotes and show very little or no copy number dynamics at all at the origin of multicellular fungi. A second category comprises gene families with a deep eukaryotic origin that show duplications coincident with the evolution of hyphae. However, no families were found that had statistically significantly elevated number of duplications in BCZ nodes, indicating limited evolutionary novelty in these nodes. In the third category there are gene families whose origin mapped to BCZ nodes (RGSs, formins, APSES and Bem1 families). Such gene families could have evolved *de novo* or diverged in sequence so much that similarity is not detectable to homologous non-fungal sequences. We find candidates for both scenarios. For example, the MedA or APSES families contain fungal-specific protein domains; these have conceivably evolved in early fungi and represent fungal-specific innovations. On the other hand, the detected formin and RGS families contained only fungal genes, but their characteristic Interpro domains occur outside of fungi too, possibly reflecting common ancestry, with evidence for homology blurred by sequence divergence.

We found that several multicellularity-related genes predate the emergence of hyphal MC and were co-opted for hyphal growth during evolution, which mirrors patterns observed in multicellular animals and plants^6,100,101^. Such observations rise to the hypothesis that in terms of genetic novelty, transitions to multicellularity represent a minor rather than a major evolutionary step^102^, an idea that finds support in the observations made here on fungi. Cell polarity maintenance, vesicle trafficking and cytoskeletal systems of unicellular eukaryotes may have turned out to be useful functions in hyphal MC, on which the extreme polarized growth of fungal hyphae could have built during evolution. The phagocytotic machinery was probably exapted for hyphal MC: its original function was most likely lifted by the emergence of a rigid cell wall in early fungi, which probably made the underlying genes dispensable. However, instead of being lost over time, its components got incorporated into hyphal MC.

Finally, beyond gene family events, certainly other genetic mechanisms (e.g. changes in amino acid sequence and domain composition, changes to biophysical properties of genes) also contributed to the evolution of hyphae. Our analyses of gene architecture revealed significant differences between MC-related genes of unicellular and multicellular fungi, indicating that changes in CDS length or intron content are also relevant for the emergence of hyphal MC^103^. Reports of sequence-level changes and domain shuffling in connection with the evolution of hyphal MC have also been published. There is evidence for fungal kinesins being 2x more processive than other eukaryotic kinesins^104^-probably in response to needs of long range transport along the hyphal axis. Similarly, class V and VII chitin synthases gained a myosin motor domain in early fungi, providing higher efficiency in polarized chitin synthesis^105–107^.

Taken together, our results suggest that multiple mechanisms probably contributed to the evolution of hyphal MC, including changes in gene structure, gene duplications, *de novo* gene family birth, and co-option/neofunctionalization. Compared to other multicellular lineages, the evolution of fungi shows several unique patterns. While the expansion of adhesion and signal transduction mechanisms is shared by most colonial and aggregative multicellular lineages examined so far^3,9,19,108–110^, we did not find evidence for this in fungi. This could be explained by the peculiar life history of fungal hyphae, which shares similarity with only Oomycota. While adhesion might not be key in vegetative hyphae, there is plenty of evidence for active communication between neighboring hyphae^111–113^. It is possible that the main modes of communication in fungal hyphae are not linked to cell surface receptors, might be related to volatiles (such as farnesol^80,114,115^) or are not known yet. These observations suggest that multicellularity in fungi differs considerably from that in other lineages and raises the possibility that hyphal MC should be considered a third, qualitatively different way in addition to the aggregative and clonal modes of evolving MC. Subjective categorizations aside, hyphal MC represents a highly successful adaptation to terrestrial life and comparative genomics opens the door for discussions on whether major phenotypic transitions represent - in terms of genetic novelty - a major or a minor transition.

## Methods

### Organismal phylogeny

We assembled a dataset containing whole proteomes of 71 species and performed all-vs-all blast using mpiBLAST 1.6.0^116^. We omitted Microsporidia from the dataset due to the high rate of evolution of this group. Proteins were clustered into gene families with Markov Cluster Algorithm (MCL)^117^ with an inflation parameter of 2.0. Clusters with at least 50% taxon occupancy were chosen and were aligned by PRANK 140603^118^ while trimAl 1.4.rev15^119^ was used to remove poorly aligned regions from the multiple sequence alignments using the parameter –gt 0.2. Approximately-maximum-likelihood gene trees were inferred by FastTree^120^ using the LG+CAT model (-lg -cat20), and the option -gamma to compute a Gamma20-based likelihoods. Using a custom-made Perl script we excluded gene trees with deep paralogs to identify single-copy genes. Alignments of single-copy gene clusters were concatenated into a supermatrix and a species tree was estimated using RAxML 8.2.4^121^ under the PROTGAMMAWAG model. The model was partitioned by gene. Bootstrapping was performed on the dataset in 100 replicates.

### Ancestral character state reconstructions

The 71 species were coded for their ability to form hyphae, either as hyphal or non-hyphal. Species that could not be unambiguously assigned to hyphal or non-hyphal (*Catenaria anguillulae*) and those with the ability to grow either as hyphae or unicells (most yeasts) were coded as uncertain. Bayesian MCMC reconstruction of ancestral character states was performed under the threshold model^122^ using Bayesian MCMC with the “ancThresh” function in phytools v0.6-60^123^ in R^124^. The number of generations for MCMC was set to 1,000,000, and the method “mcmc” was used with the Brownian motion as the model for the evolution of the liability. Burn-in parameter was set to default. Convergence was checked by inspecting likelihood values through time.

### Analyses of Gene family evolution

To investigate the evolutionary dynamics of gene families containing hyphal morphogenesis-related genes, we performed all-vs-all blast (NCBI Blast 2.7.1+)^125^ for proteomes of the 71 species and did sequence similarity-based protein clustering by following the MCL clustering protocol^117^ used by Ohm et al 2012^126^. The resulting protein clusters were aligned by PRANK 140603^118^ with default parameters, and ambiguously aligned regions were removed using trimAl 1.4.rev15^119^ with the argument –gt 0.2. MAFFT v7.222^127^ (option –-auto) was used as an alternative alignment tool for clusters that could not be aligned by PRANK due to computational limitations (80 out of 34032 clusters). Maximum Likelihood inference of gene trees and calculation of Shimodaira-Hasegawa-like branch support values were carried out in RAxML 8.2.4^128^ under the PROTGAMMAWAG model of protein evolution. The calculated SH-like branch support values were used in gene tree-species tree reconciliation in Notung-2.9^129^. An edge-weight threshold of 0.9 was used, as SH-like support values are usually less conservative than ML bootstrap values (where 70% is usually taken as indication of strong support). Reconciliation was performed on the maximum likelihood gene trees and the ML species tree for the 71 species as input. We reconstructed the gene duplication/loss dynamics of gene families along the species tree using respective scripts from the COMPARE pipeline^94,130^. The numbers of gains and losses for each gene family and for each branch of the species tree were recorded and mapped on the species tree. Ancestral gene copy numbers were calculated for every internal node, summing the mapped duplications and losses across the species tree. Mappings were generated for each of the functional groups and also for kinases, adhesion-related proteins, receptors as well as for all gene families across the 71 genomes.

To test if genes related to hyphal MC experience an episode of increased duplication rate in nodes where hyphal growth putatively originated (BCZ nodes), we performed gene duplication enrichment analysis for each of the 362 families and for functional groups. To test if a cluster or a functional group shows significantly more or less duplications than expected by chance in BCZ nodes, we run two-tailed Fisher’s exact tests (p < 0.05). We compared the number of duplications mapped to BCZ nodes for a given gene family to the genome-wide number of duplications in BCZ nodes, using total number of duplications across the tree as a reference.

### Analyses of key multicellularity-related genes

The above strategy was used to reconstruct the evolution of kinase, receptor and adhesion-related gene families. Protein clusters containing kinase genes, both serine-threonine kinases and histidine kinases, were collected based on InterPro domains. Identification of serine-threonine kinases and histidine kinases followed Park et al 2011^131^ and Herivaux et al 2016^88^, respectively. Classification of histidine kinases followed Defosse et al^87^.

Families of adhesive proteins were identified based on experimentally characterized genes collected from the literature. We identified 45 genes, which mostly grouped into flocculins, lectins, hydrophobins and other (GPI)-modified cell wall adhesins. We identified receptor genes based on InterPro domains that are annotated with the gene ontology term ‘receptor activity’ (but not ‘receptor binding’ or other terms indicative of indirect relationships to receptor function), resulting in 27 IPR terms (IPR000161, IPR000276, IPR000337, IPR000363, IPR000366, IPR000481, IPR000832, IPR000848, IPR001103, IPR001105, IPR001499, IPR001546, IPR001946, IPR002011, IPR002185, IPR002280, IPR002455, IPR002456, IPR003110, IPR003292, IPR003980, IPR003982, IPR005386, IPR006211, IPR017978, IPR017979, IPR017981).

### Analyses of phagocytosis-related genes

We collected information on phagocytosis-related genes from recent reviews on *Dictyostelium*^65,66^, identified the corresponding genes of this species in our dataset and the protein clusters that contained homologs of the identified genes. Mapping of gene duplications and losses along the species tree was done as described above.

### Genome-wide screen for putative hyphal multicellularity-related gene families

To identify gene families with increased rates of gene duplication coinciding with the origin of hyphal MC, we set up a pipeline that tests for higher than expected rate of duplication in nodes of the species tree to which the origin of hyphae could be located (BCZ nodes). For each gene family, gene duplication rates in BCZ nodes were compared to duplication rates of the same family in other parts of the species tree (nodes before and after BCZ nodes). Gene duplication rates were computed by dividing the number of reconstructed duplications for a given branch by the length of that branch using a custom Perl script. Terminal duplications and duplications mapped to metazoan ancestors were excluded from the analysis. The resulting node×duplication rate matrix was analyzed by a two-factor permutation ANOVA^132^ with degrees of freedom DFT=2, in R, with P < 0.05 considered as significant. We further required that the detected clusters be conserved (>=1 copy) in at least 70% of filamentous fungi.

### Analyses of gene losses in yeast-like fungi

We analyzed gene losses in five yeast-like lineages by comparing the number of losses genome-wide, to the numbers of losses in hyphal morphogenesis related genes (actin cytoskeleton regulation, polarity maintenance, cell wall biogenesis/remodelling, septation and septal plugging, signal transduction, transcriptional regulation, vesicle transport, microtubule-based transport and cell cycle regulation) relative to ancestral copy numbers. P-values were calculated by Fisher’s exact test, with P < 0.05 considered as significant. The percentage of genes retained in yeast genomes was calculated for every functional category by comparing ancient gene-copy number prior to the emergence of yeast-like lineages to the average gene copy-number of terminals.

### Statistical analysis of genomic features

An R script (available upon request) was written to generate coding sequence (CDS)/intron statistics (strand, order, length, count) based on genome annotations of the 71 species. CDS feature coordinates for each gene were extracted and subsequently used to calculate intron coordinates. Statistical significance of the differences between the gene, CDS and intron lengths of 4 unicellular and 39 multicellular fungi was investigated by independent two-tailed Welch’s t-test with pooled variance estimation (var.equal=FALSE), using the t.test function in R.

## Data availability

Files associated with this paper (including gene and species trees, gene duplication/loss catalogs) have been deposited in Dryad (Accession number, *to be provided upon publication*).

## Acknowledgements

The authors are thankful to Sándor Kocsubé for his help in crafting figures. This work was supported by the ‘Momentum’ program of the Hungarian Academy of Sciences (contract No. LP2014/12 to L.G.N.) and the European Research Council (grant no. 758161 to L.G.N.). E.K. acknowledges support from the National Talent Program of the Ministry of Human Capacities (contract No. NTP-NFTÖ-16-0566) and the Young Researchers Program of the Hungarian Academy of Sciences. N.T. acknowledges support from Japan Society for the Promotion of Science (JSPS) KAKENHI (grant number: 18K05545) and Japan Science and Technology Agency (JST) ERATO (grant number: JPMJER1502).

## Author contributions

K. and N.G.L. concieved the study. E.K. collected literature data on morphogenesis-related genes. E.K. and A.N.P. inferred species trees, B.B. performed clustering, E.K., K.K., T.K., Z.M. and T.V. analyzed gene family evolution. B.H. analyzed gene structural changes. E.K., M.R., N.T. and N.G.L. interpreted the results and wrote the paper. All authors have read and commented on the manuscript.

## Supplementary Figures

**Supplementary Figure 1.**
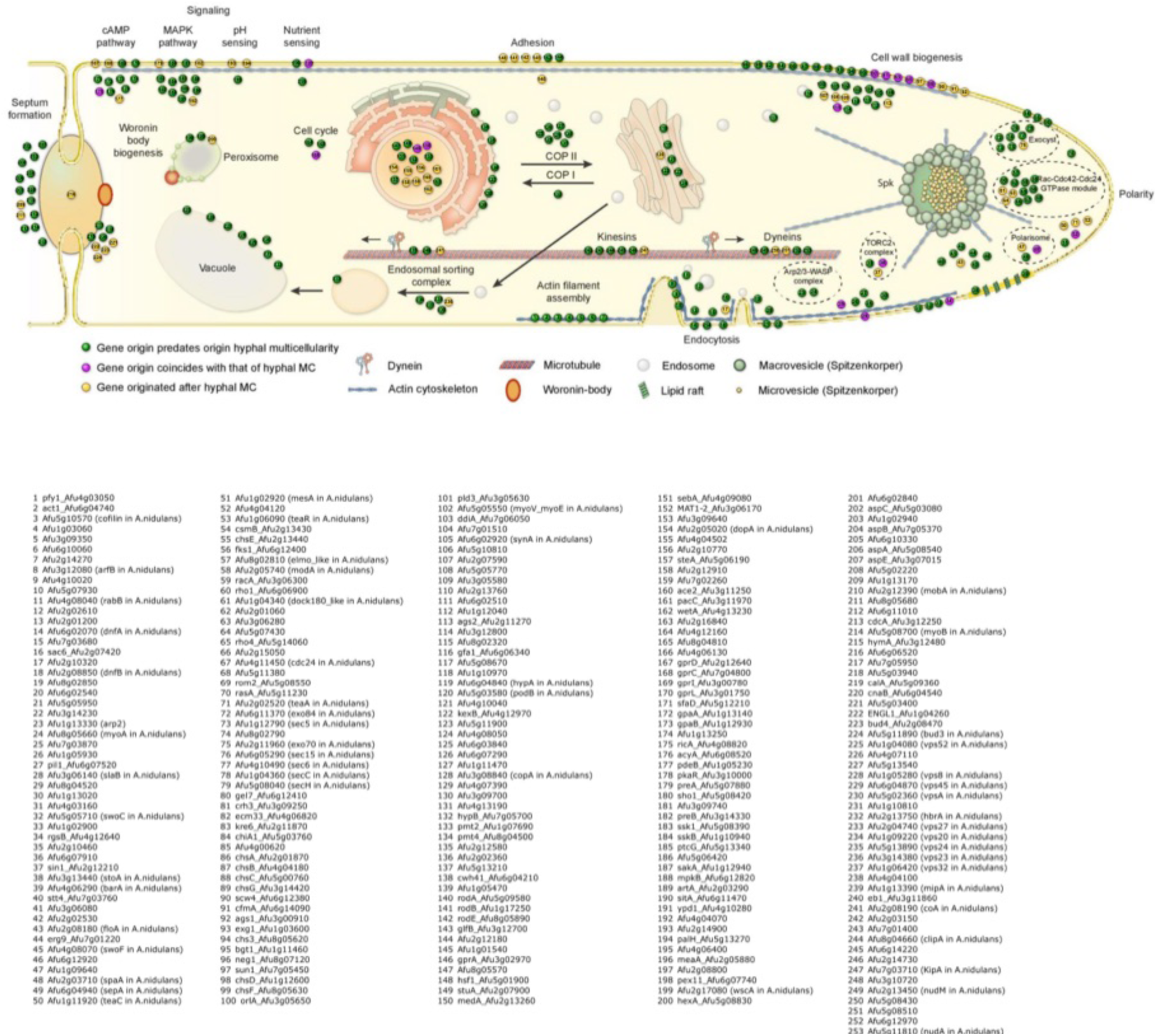
Phylogenetic age distribution of hypha morphogenesis genes. Figure mirrors main text Figure 2 with gene names provided for each dot.

**Supplementary Figure 2.**
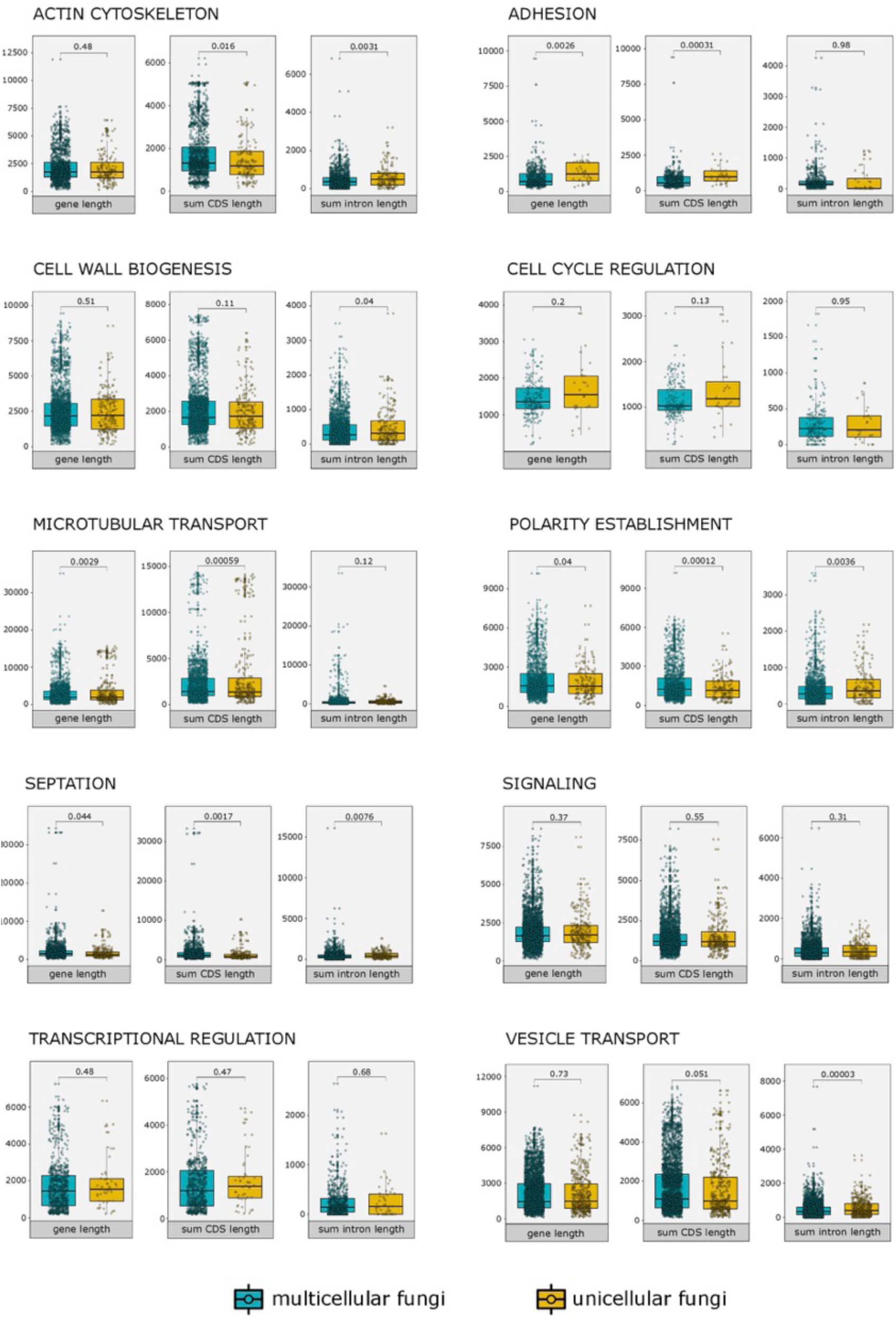
Statistical comparisons of basic structural properties of genes related to hyphal MC. Box plots display differences in gene, CDS and intron lengths of 4 unicellular (yellow) and 39 multicellular fungi (blue) in nine functional categories.

**Supplementary Figure 3.**
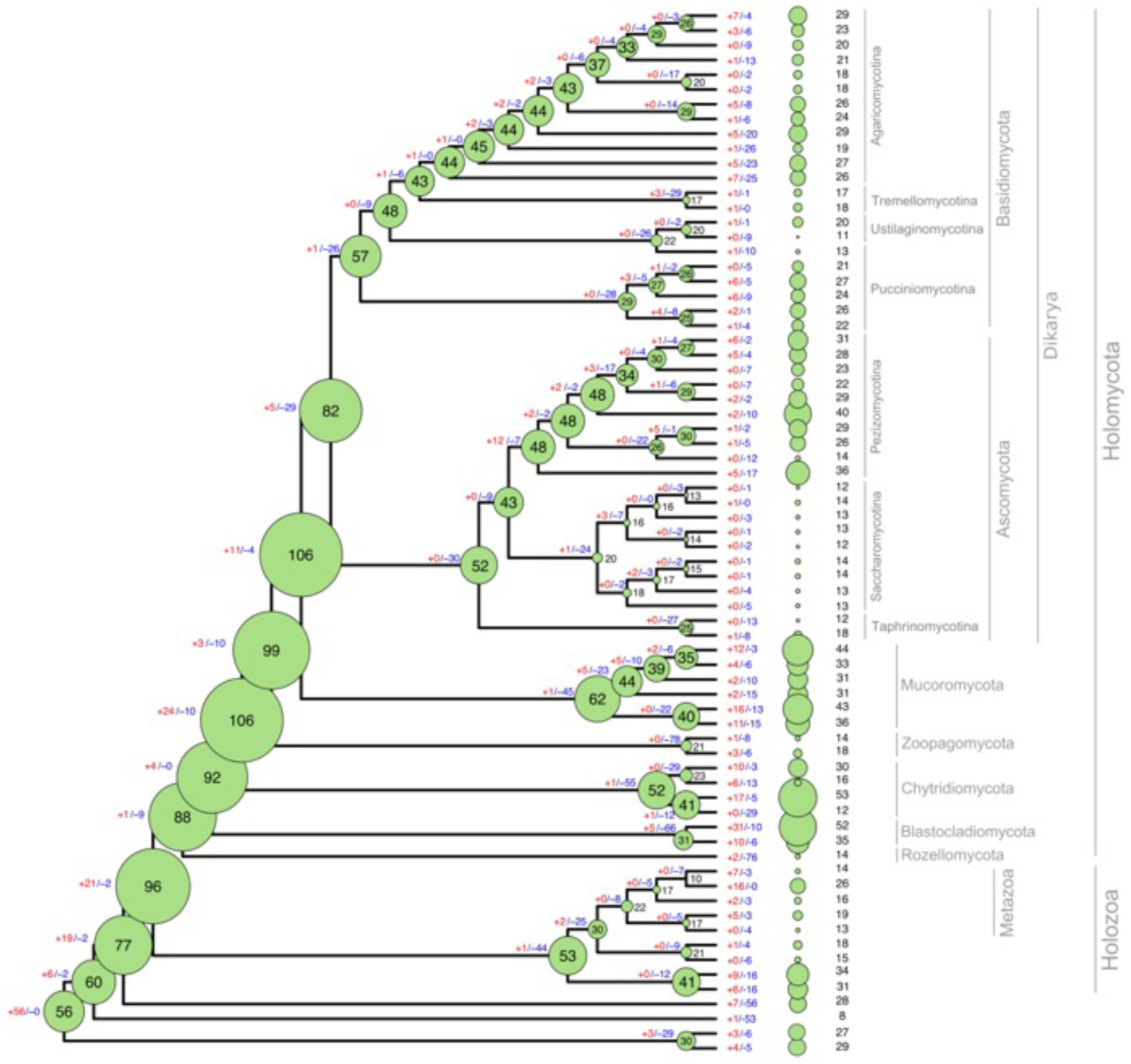
Reconstructions of ancestral gene copy numbers of histidine-kinase genes. Numbers at the branches represent gene duplications (+) and losses (-) inferred by COMPARE. Bubble size is proportional to reconstructed ancestral gene copy number. Copy number distribution for each species is shown right to the tree.

**Supplementary Fig. 4.**
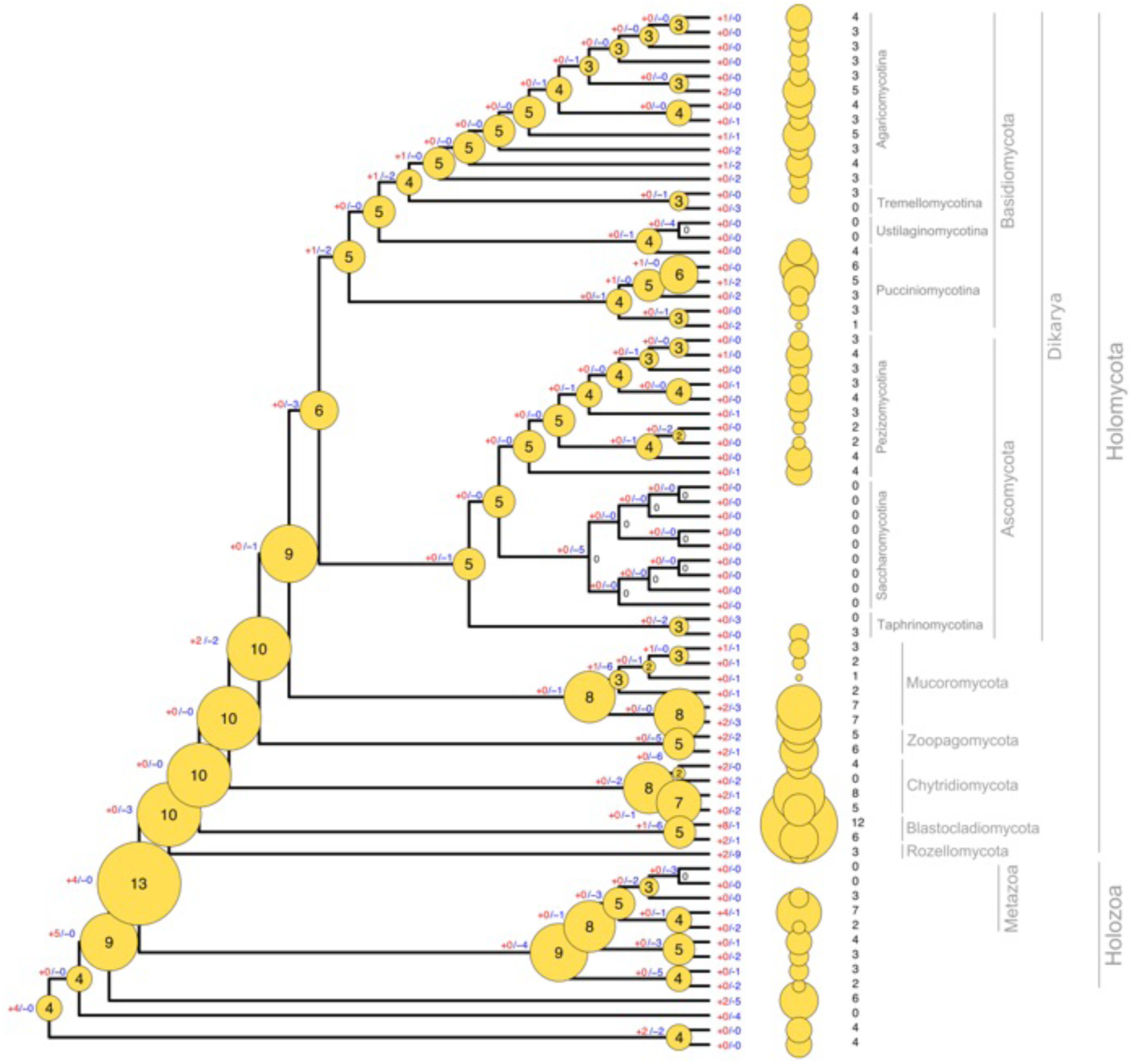
Reconstructions of ancestral gene copy numbers of NADPH-oxidase genes. Numbers at the branches represent gene duplications (+) and losses (-) inferred by COMPARE. Bubble size is proportional to reconstructed ancestral gene copy number. Copy number distribution for each species is shown right to the tree.

